# Acid Experimental Evolution of the Extremely Halophilic Archaeon *Halobacterium* sp. NRC-1 Selects Mutations Affecting Arginine Transport and Catabolism

**DOI:** 10.1101/662882

**Authors:** Karina S. Kunka, Jessie M. Griffith, Chase Holdener, Katarina M. Bischof, Haofan Li, Priya DasSarma, Shiladitya DasSarma, Joan L. Slonczewski

**Author notes:** Corresponding Author: Joan L. Slonczewski, Department of Biology, Kenyon College, Gambier, Ohio, USA.

## Abstract

**Background:** *Halobacterium* sp. NRC-1 (NRC-1) is an extremely halophilic archaeon that is adapted to multiple stressors such as UV, ionizing radiation and arsenic exposure. We conducted experimental evolution of NRC-1 under acid stress. NRC-1 was serially cultured in CM+ medium modified by four conditions: optimal pH (pH 7.5), acid stress (pH 6.3), iron amendment (600 μM ferrous sulfate, pH 7.5), and acid plus iron (pH 6.3, with 600 μM ferrous sulfate). For each condition, four independent lineages of evolving populations were propagated. After 500 generations, 16 clones were isolated for phenotypic characterization and genomic sequencing.

**Results:** Genome sequences of all 16 clones revealed 378 mutations, of which 90% were haloarchaeal insertion sequences (ISH) and ISH-mediated large deletions. This proportion of ISH events in NRC-1 was five-fold greater than that reported for comparable evolution of *E. coli*. One acid-evolved clone had increased fitness compared to the ancestral strain when cultured at low pH. Seven of eight acid-evolved clones had a mutation within or upstream of *arcD*, which encodes an arginine-ornithine antiporter; no non-acid adapted strains had *arcD* mutations. Mutations also affected the *arcR* regulator of arginine catabolism, which protects bacteria from acid stress by release of ammonia. Two acid-adapted strains shared a common mutation in *bop*, which encodes the bacteriorhodopsin light-driven proton pump. Unrelated to pH, one NRC-1 minichromosome (megaplasmid) pNRC100 had increased copy number, and we observed several mutations that eliminate gas vesicles and arsenic resistance. Thus, in the haloarchaeon NRC-1, as in bacteria, pH adaptation was associated with genes involved in arginine catabolism and proton transport.

**Conclusions:** Our study is among the first to report experimental evolution with multiple resequenced genomes of an archaeon. Haloarchaea are polyextremophiles capable of growth under environmental conditions such as concentrated NaCl and desiccation, but little is known about pH stress. *Halobacterium* sp. NRC-1 (NRC-1) is considered a model organism for the feasibility of microbial life in iron-rich brine on Mars. Interesting parallels appear between the molecular basis of pH adaptation in NRC-1 and in bacteria, particularly the acid-responsive arginine-ornithine system found in oral streptococci.

## INTRODUCTION

*Halobacterium* sp. NRC-1 (NRC-1) is a polyextremophile that grows optimally at NaCl concentrations in excess of 4 molar (1). A genetically tractable model microbe (2), it was the first halophilic Archaeon with a fully sequenced genome (3). Besides high salt, NRC-1 is capable of surviving: high doses of ionizing radiation and dessication (4), UV radiation (5), temperature extremes (6), and toxic ions such as arsenite (7). These traits have made NRC-1 a model for studying the possibility of life outside Earth under conditions such as the stratosphere (8,9) or on Mars (10–12).

Water on Mars contains high concentrations of salt, as well as acid and iron (13). The Mars Exploration Rover Opportunity discovered substantial deposits of an iron hydrous sulfate mineral known as jarosite [KFe^3+^_3_(OH)_6_(SO_4_)_2_] which forms in acidic and iron-rich aqueous environments. On earth such conditions occur in acid mine drainage and near volcanic vents. Opportunity’s discovery of jarosite on Mars was evidence of acidic, liquid water and an oxidizing atmosphere in the Martian past (13,14). Occuring together, acid and metals can amplify the stress associated with each condition (15). Thus, it is of interest to investigate how a neutralophilic halophile such as NRC-1 (16) might adapt to conditions of acid and high iron.

An informative approach to examine the genomic basis of stress response is experimental laboratory evolution (17–23). Experimental evolution of bacteria reveals changes in phenotype and genotype in response to specific stressors in a controlled environment, such as carbon source limitation or extreme pH. In bacterial adaptation to various kinds of pH stress, we find a recurring pattern that dominant responses to short-term stress actually decrease fitness over many generations of long-term exposure. For example, amino-acid transport and catabolism play important roles in extreme-acid survival of *Escherichia coli* (24,25). However, 2,000 generations of *E. coli* evolution at pH 4.8 select for loss of three acid-inducible amino-acid decarboxylase systems, including arginine decarboxylase (21). As a membrane-permeant acid, benzoic acid induces glutamate decarboxylase and drug resistance regulons, yet these systems are lost or downregulated during experimental evolution (26),(20). At high external pH, *E. coli* survival requires the stress sigma factor RpoS; however, generations of growth at high pH select against RpoS expression and activity (27). It is therefore of interest to investigate whether similar patterns of reversal occur in archaea.

Relatively few experimental evolution studies have been reported in archaea. In NRC-1, serial application of lethal doses of ionizing radiation selected more resistant mutants that had increased expression of a single-strand DNA binding protein (28). In the thermoacidophile *Sulfolobus solfatericus*, serial passage in extreme acid yielded strains that grow below pH 1 (29). These strains showed mutations in amino acid transporters, as well as upregulation of membrane biosynthesis and oxidative stress response. In *Metallosphaera sedula*, serial passage led to a pH 0.9-adapted strain with four mutations, one of which is an amino-acid/polyamine transporter (30). These findings are intriguing, given the role of amino-acid transport and catabolism in extreme-acid survival of bacteria (24,25). For example, arginine transport and catabolism, which yields CO_2_ plus two ammonium ions, is a prominent response to acid stress of oral streptococci (31),(32).

Archaea employ various processes that involve proton transport via primary pumps and antiporters (24,33,34). *Halobacterium* strains possess the light-driven proton pump bacteriorhodopsin (*bop*) that generates proton motive force (PMF) (35,36) as well as several sodium-proton antiporters, which export sodium in exchange for protons (6).

We conducted experimental evolution of NRC-1 under conditions of low pH (pH 6.5-6.3) and at optimal pH for growth (pH 7.5), with high iron versus low iron concentration. The NRC-1 genome includes a main chromosome and two minichromosomes or megaplasmids (3,37). It accumulates frequent IS mutations (38,39) which may mediate rapid adaptations to environmental stress. Our study of experimental evolution in a haloarchaeon assesses which mutations contribute to archaeal evolution in acid stress. Here we describe analysis of phenotypic changes across evolved clones from each population, and then use genomic analysis to identify potential underlying mutational bases of these phenotypic responses to selection. Genome analysis of 16 clones revealed a remarkable proportion of events mediated by insertion sequences (ISH). In acid-adapted strains, we found a high frequency of mutations in the arginine-ornithine antiporter *arcD* (40) and in the associated *arcR* arginine catabolism regulator (41).

## RESULTS

### Experimental evolution under conditions of acid and iron stress

Serial culture of evolving populations was conducted as described under Methods (Additional File 1, Fig. S1). Populations of NRC-1 were founded from a single clone and cultured in modified CM^+^ medium (2,3) with appropriate buffers to maintain pH. Each population was diluted 500-fold every four days (approximately 9 generations). Four independent populations were maintained for each condition: the optimal growth condition, pH 7.5 (designation M); acid stress, initially pH 6.5, later pH 6.3 (designated J); iron amendment, pH 7.5 with 600 µM ferrous sulfate (designated S); and acid with iron amendment (designated K) for a total of 16 experimental populations. Populations evolved under acid stress were cultured at an initial pH of 6.5, which was then lowered to 6.3 at generation 250, as the populations adapted.

After all populations reached 500 doublings, two clones were isolated from each population by three rounds of streaking on CM+ agar for a total of 32 evolved clones. Genomic DNA was extracted from 16 of these clones, and from the founder stock of NRC-1. DNA samples were sequenced by Illumina MiSeq, and mutations were identified by comparison of the “evolved strain” sequences to that of the NRC-1 ancestral stock, assembled on the reference genome (3) using the *breseq* pipeline (42–44). The strains we characterized are listed in **Table 1**.

**Table 1.**
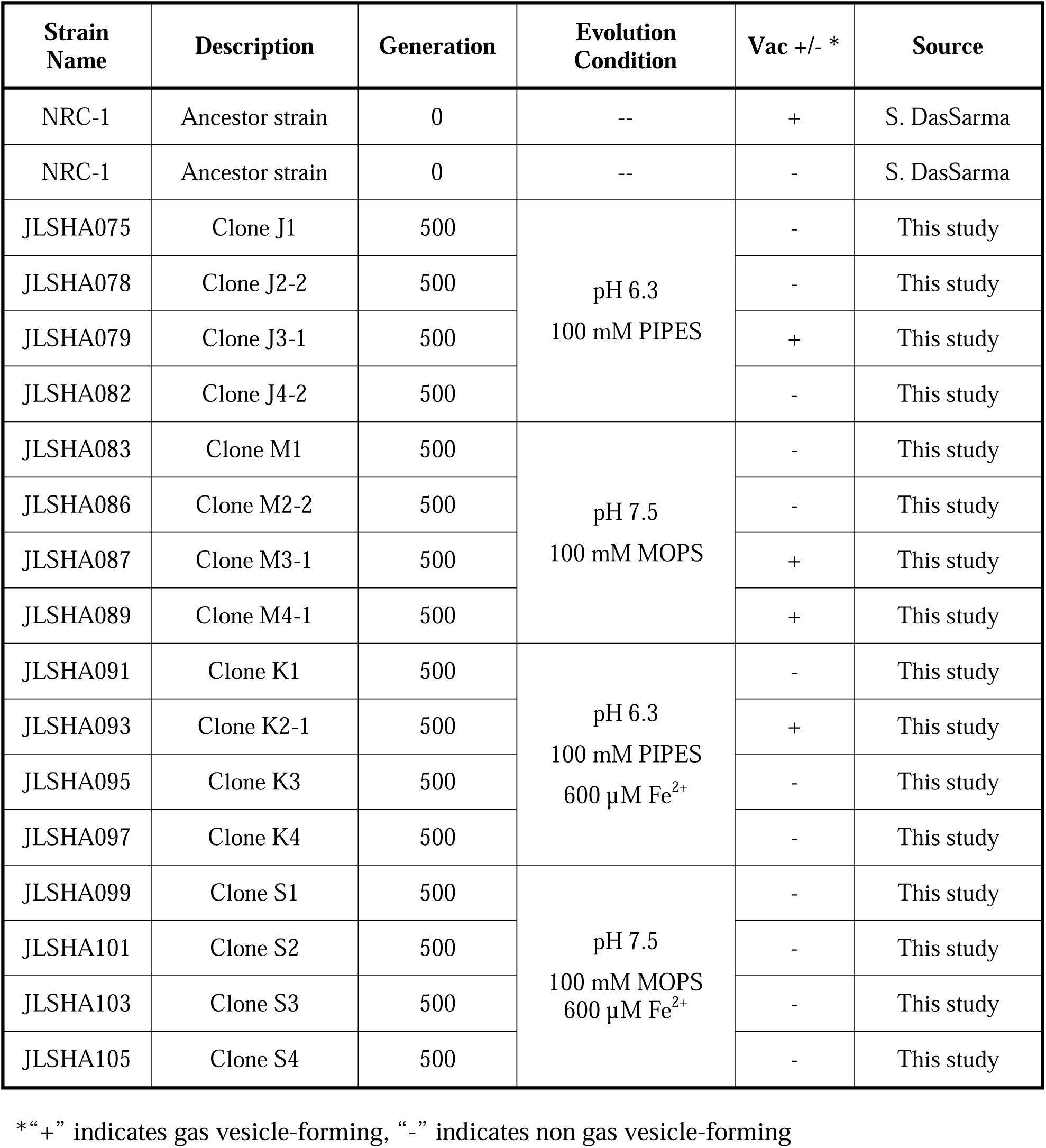
Strains used in this study

### Mutations in the genomes from evolving populations

The genomes of the 16 clones were compared to those of the NRC-1 ancestor which we resequenced from our lab stock (**Tables S1, S2, S3**). The genome of our NRC-1 stock was also compared to that of the NCBI reference sequence *Halobacterium* sp. NRC-1 (3) as shown in Additional File 1, **Table S4**. A small number of positions differed from that of the reference. Some of these differences are consistent with those of later sequence reports (45,46). The sequences differences shown in **Table S4** were excluded in our analysis of the evolved clones.

The genomes of the evolved clones had a total of 378 mutations, of which 349 were unique to one strain at the base-pair level. Representative mutations of interest are summarized in **Table 2**. Mutation frequencies were compared for the main replicon and minichromosomes. In total across all resequenced genomes there were 120 mutations in minichromosome pNRC100, and 171 mutations on minichromosome pNRC200. pNRC100 is about 10% as long as the main chromosome, and pNRC200 is about 20% as long; thus, the two minichromosomes had a mutation frequency more than ten-fold greater than that of the main chromosome, a finding consistent with previous reports of plasmid or minichromosome mutation (3).

**Table 2.**
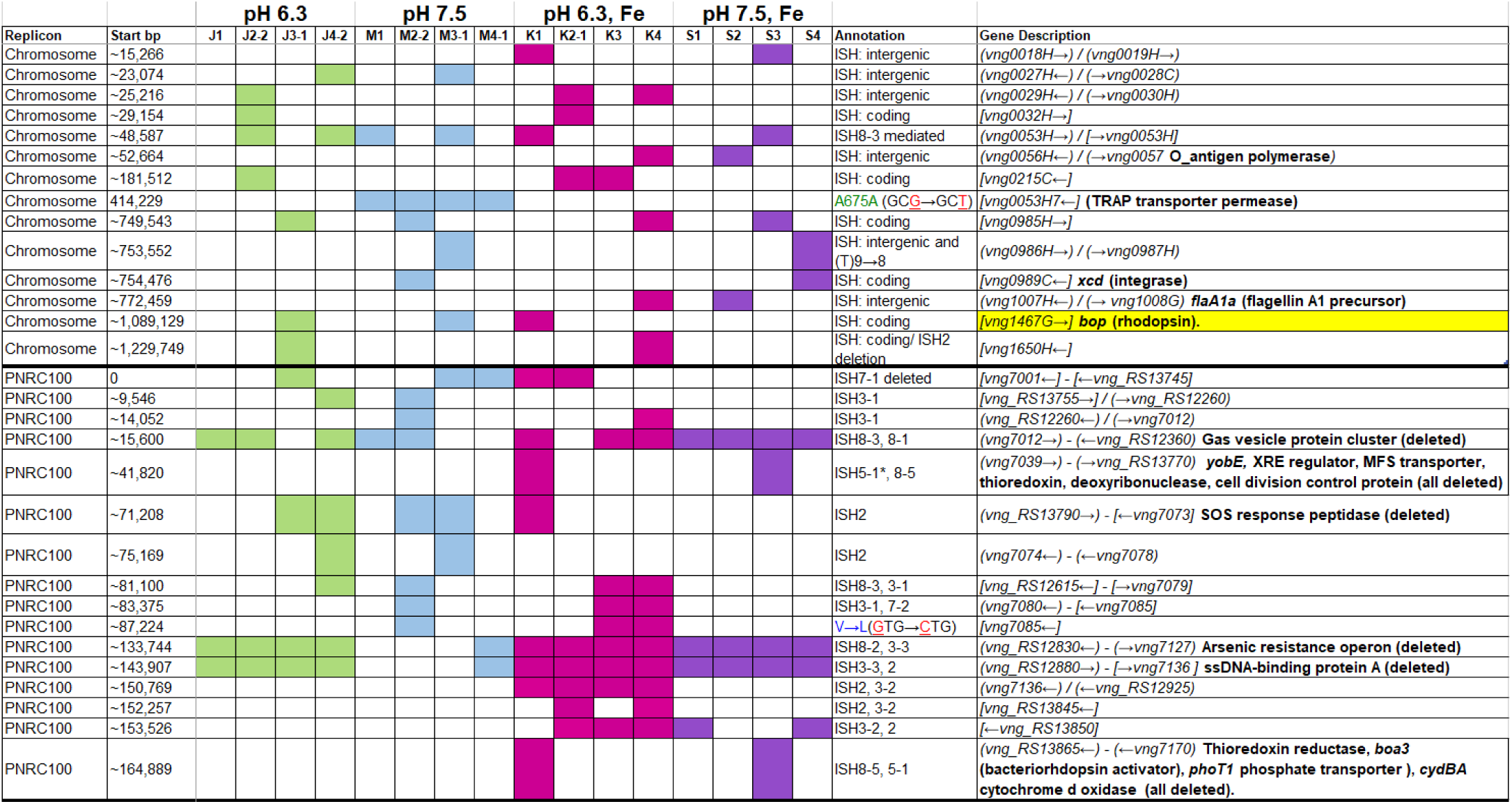
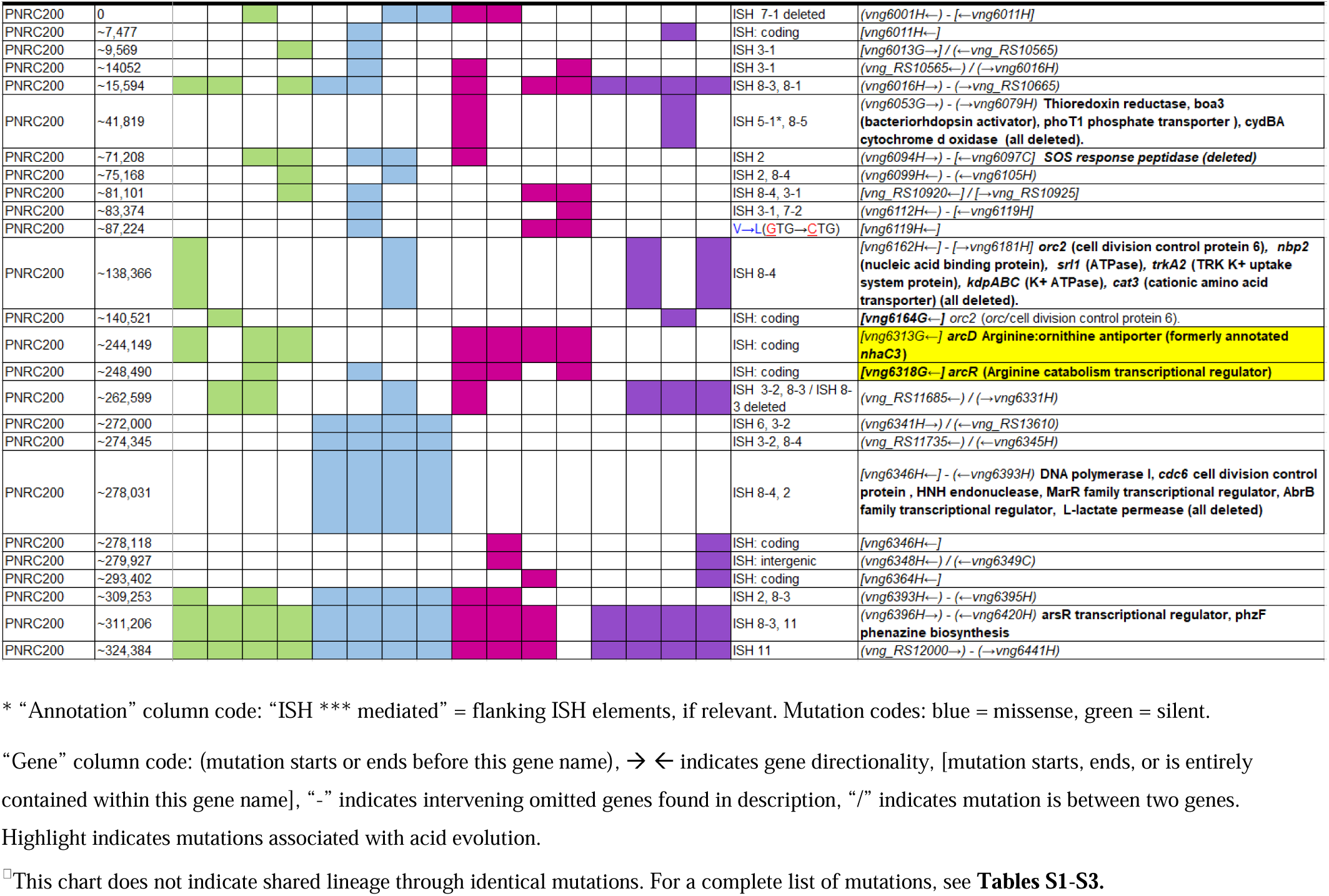
Selected mutations found in evolved clones.*^□^

In the 16 clones, overall, 87 different mutations were found on the main chromosome. Of these, 90% consisted of new ISH positions, or of deletions mobilized by existing ISH elements (**Table 3**). Mutation distributions of the minichromosome replicons showed no significant difference in ISH proportion (94 mutations out of 111, on pNRC100; 141 out of 156 on pNRC200). For comparison, we considered a recent *breseq* analysis of 16 *E. coli* genomes following 500 generations evolution with an organic acid (26). Only 18% of the *E. coli* mutations were mediated by insertion sequences. Thus, NRC-1 evolution showed five-fold greater proportion of insertion sequence activity than *E. coli*. Our quantitative analysis of experimentally evolved genomes is consistent with earlier evidence of high ISH activity in halobacterial genomes (38),(47–50),(45).

**Table 3.**
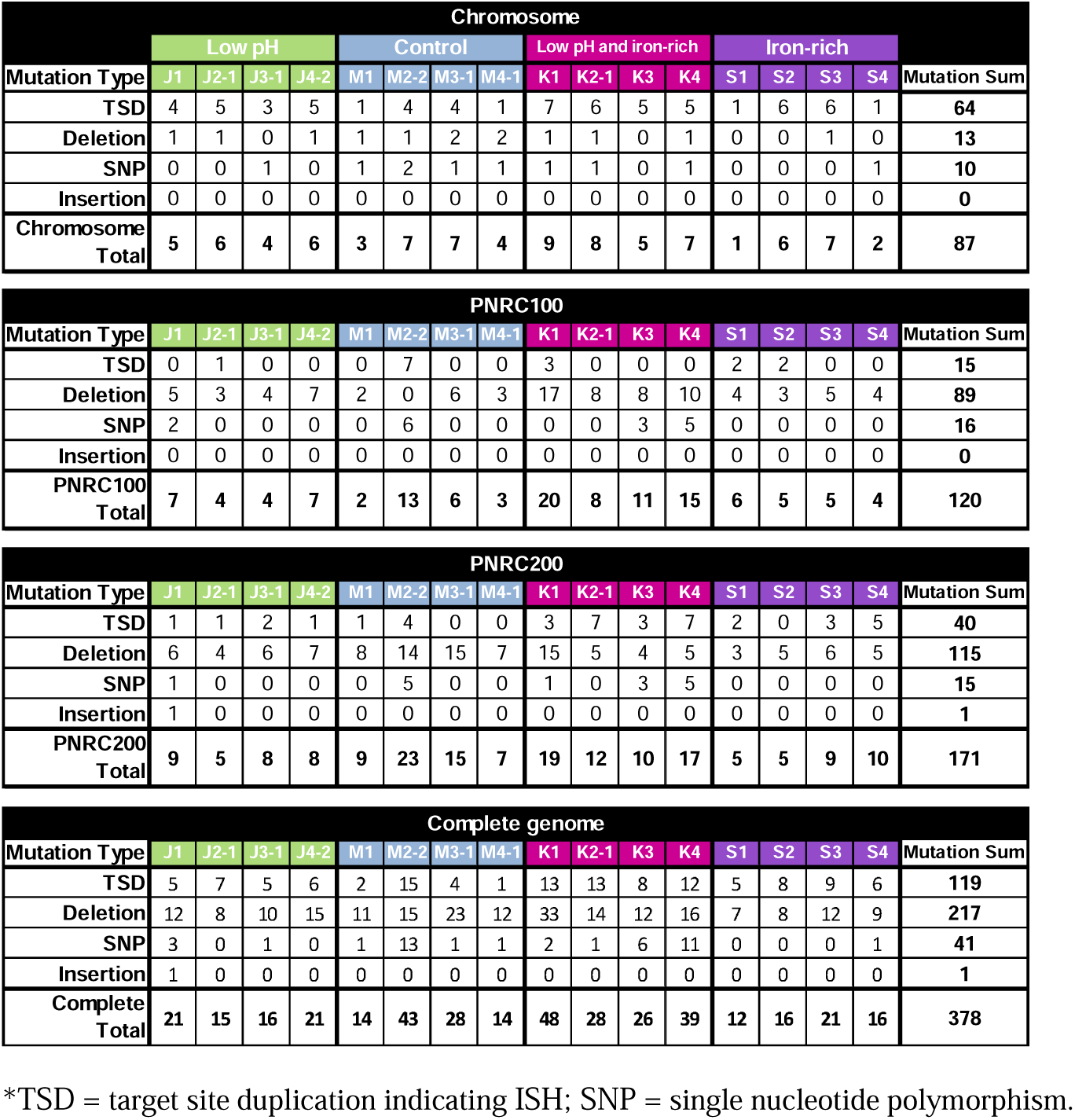
Classes of mutations found in evolved clones.*

Haloarchaea including *Halobacterium salinarum* species are known for polyploidy (15-25 genome copies per cell) and for ploidy variation among replicons within a cell (51). Our evolved clones showed evidence for variable ploidy between and within replicons. Mean read coverage by replicon was modeled by *breseq* (**Table 4**).

**Table 4.**
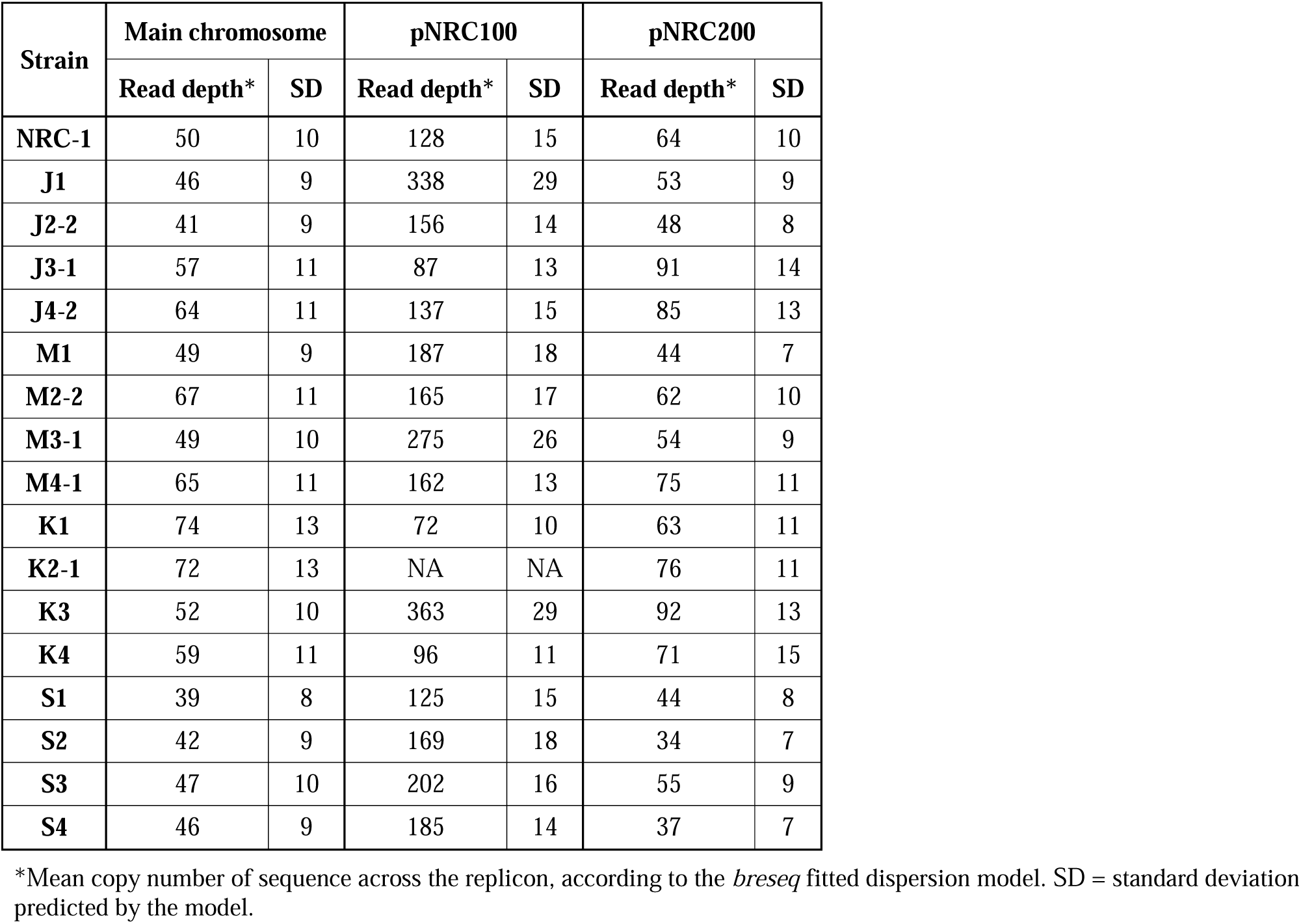
Coverage depth for NRC-1 and evolved clones.

| **Strain** | **Main chromosome** |  | **pNRC100** |  | **pNRC200** |  |
| --- | --- | --- | --- | --- | --- | --- |
|  | **Read depth\*** | **SD** | **Read depth\*** | **SD** | **Read depth\*** | **SD** |
| **NRC-1** | 50 | 10 | 128 | 15 | 64 | 10 |
| **J1** | 46 | 9 | 338 | 29 | 53 | 9 |
| **J2-2** | 41 | 9 | 156 | 14 | 48 | 8 |
| **J3-1** | 57 | 11 | 87 | 13 | 91 | 14 |
| **J4-2** | 64 | 11 | 137 | 15 | 85 | 13 |
| **M1** | 49 | 9 | 187 | 18 | 44 | 7 |
| **M2-2** | 67 | 11 | 165 | 17 | 62 | 10 |
| **M3-1** | 49 | 10 | 275 | 26 | 54 | 9 |
| **M4-1** | 65 | 11 | 162 | 13 | 75 | 11 |
| **K1** | 74 | 13 | 72 | 10 | 63 | 11 |
| **K2-1** | 72 | 13 | NA | NA | 76 | 11 |
| **K3** | 52 | 10 | 363 | 29 | 92 | 13 |
| **K4** | 59 | 11 | 96 | 11 | 71 | 15 |
| **S1** | 39 | 8 | 125 | 15 | 44 | 8 |
| **S2** | 42 | 9 | 169 | 18 | 34 | 7 |
| **S3** | 47 | 10 | 202 | 16 | 55 | 9 |
| **S4** | 46 | 9 | 185 | 14 | 37 | 7 |

\*Mean copy number of sequence across the replicon, according to the *breseq* fitted dispersion model. SD = standard deviation predicted by the model.

Overall, within the ancestor and the evolved clones, the read coverage for the main chromosome was consistent with that of the minichromosome pNRC200. However, the mean coverage of the shorter minichromosome pNRC100 (191 kb) was more than twice that of the main chromosome, for our ancestral NRC-1 and for 12 of the 16 evolved clones. Clones J1, M3-1, K3, S2, and S3 had mean coverage of pNRC100 more than four-fold greater than that of the main chromosome. These high coverage ratios could indicate that our original NRC-1 stock has a double copy number of minichromosome pNRC100, relative to the main chromosome; and that some descendant clones have increased relative copy number. However, the calculations are complicated by wide variation in read coverage between different segments of the same replicon, especially in pNRC100. This variation in read coverage may be caused by internal repeats within the replicon (35). Interpretation of the data is complicated by the presence of massive deletions (Additional File 1, **Table S2**) which comprise up to 50% of the ancestral sequence (for example in clone K1) (50). Variation in read coverage could indicate the presence of plasmid copies with different deletion levels within a given polyploid cell.

### Acid-evolved clone J3-1 has a growth advantage over a range of pH values

After 500 generations of serial culture under four conditions, clones were isolated from the evolving populations. The clones were tested for genetic adaptation under various growth conditions. Each evolved clone was cultured in parallel with the ancestral strain NRC-1. The loss of gas vesicles (Vac^-^ phenotype) alters their OD_600_ reading (38,47); for this reason, clones that had lost gas vesicles were cultured in parallel with a Vac^-^ isolate of NRC-1 ancestor.

The growth of acid-evolved J-population clones was compared to that of the NRC-1 ancestor (Vac^+^) (**Figs. 1 and 2**). Clone J3-1 reached a significant two-fold higher culture density than did the ancestor when cultured at pH 6.1 or at 6.3 (**Fig. 1B**). Growth advantage was seen for all four replicate cultures of J3-1 at pH 6.1 and at pH 6.3, whereas the difference from NRC-1 cultures disappeared at pH 7.2 and at pH 7.5. Thus, strain J3-1 exhibits an acid-specific fitness advantage. The other acid-evolved J-population strains, however, had no significant growth advantage compared to NRC-1, under the conditions tested (**Fig. 2**).

**Figure 1.**
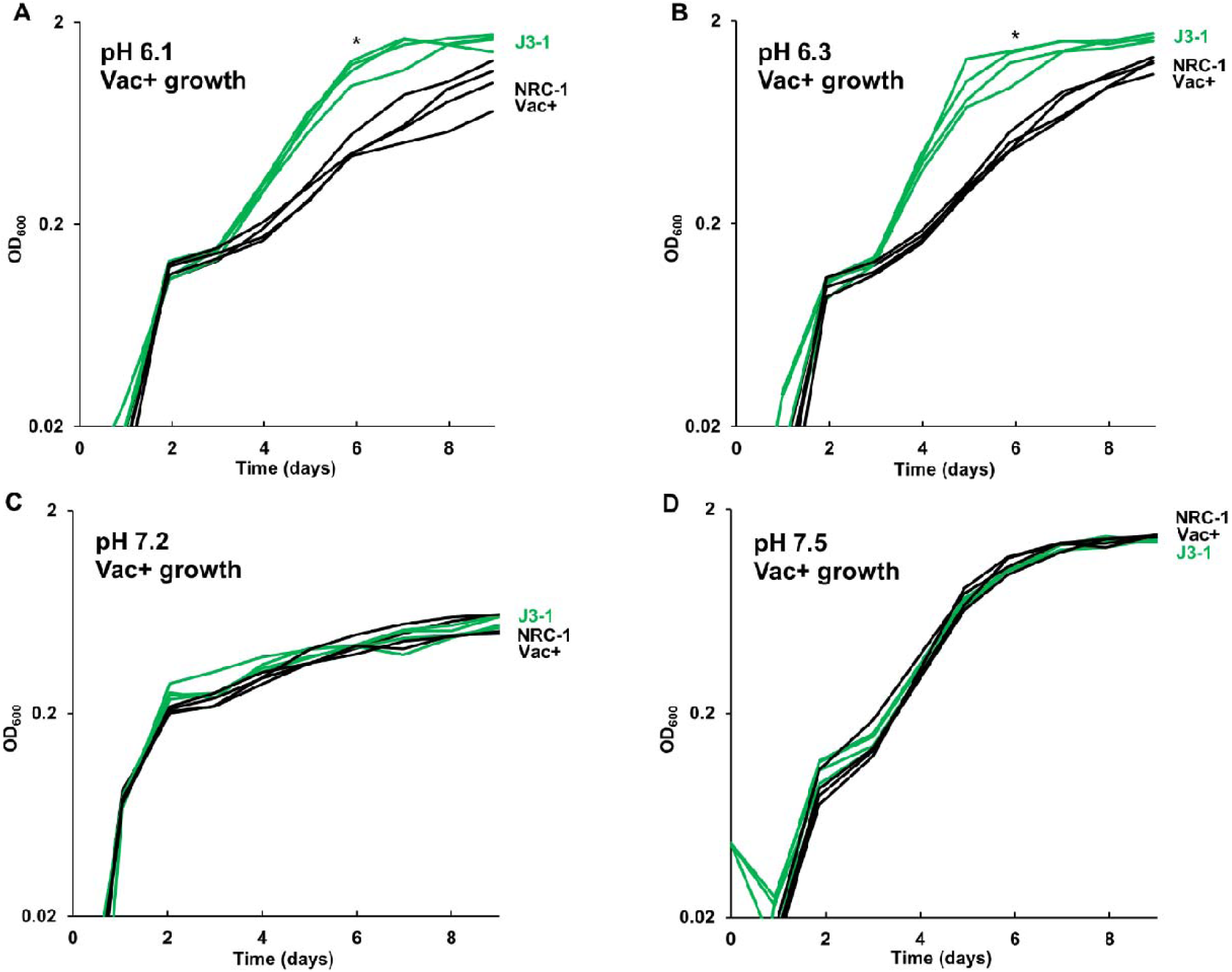
Acid-evolved clone J3-1 shows a pH-dependent growth rate increase compared to NRC-1. Growth medium consisted of CM^+^ buffered at (**A**) pH 6.1 with 100 mM PIPES; (**B**) pH 6.3 with 100 mM PIPES; (**C**) pH 7.2 with 100 mM MOPS; or (**D**) pH 7.5 with 100 mM MOPS. Representative curves of three replicates are shown. For J3-1 and NRC-1, the OD_600_ values at 144 h were compared by two-tailed t-test. At pH 6.1, P = 0.002; at pH 6.3, P = 0.01; at pH 7.2, P = 0.91; at pH 7.5, P = 0.45. “*” indicates significant endpoint growth increase from NRC-1 ancestor in at least 2 replicates.

**Figure 2.**
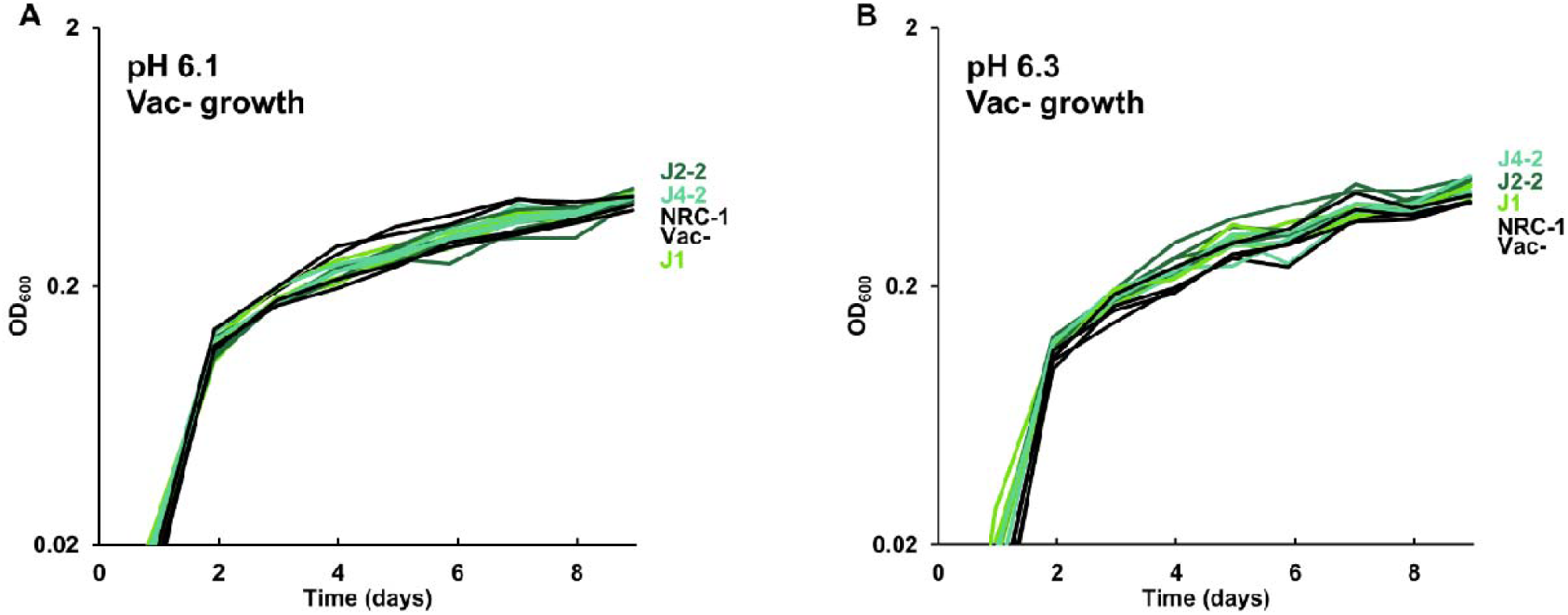
Growth of acid-evolved clones J1, J2-2, J4-2. Growth medium consisted of CM^+^ pH 6.3 with 100 mM PIPES, at (**A**) pH 6.1, (**B**) pH 6.3. Cultures were diluted from a 7-day culture in CM^+^ pH 7.2. Gas vesicle-deficient clones were compared to gas vesicle-deficient ancestral mutant NRC-1 and cell density values post log-phase (OD_600_ at 6 days) were analyzed using ANOVA with Tukey post-hoc. Representative curves of three replicates are shown.

### Acid-adapted clones shared mutations in *arcD* and in *arcR*

We inspected the genomes of acid-adapted populations J and K (acid with iron supplement) for mutations in specific genes that were not found in the populations evolved at pH 7.5. Seven out of eight of the J and K clones (but no M or S clones) had ISH mutations in or upstream of gene VNG_6313G (**Table 2**). This gene was originally classified as encoding a sodium-proton antiporter (*nhaC3*) but was shown instead by physiological experiments to encode an arginine-ornithine antiporter ArcD (52). PCR amplification and Sanger sequencing of the mutant *arcD* alleles confirmed the presence of insertion sequences ISH2 (strains K1 and K4) and ISH4 (strains J1, K2-1, K3) (**Table 5**; Additional File 1, **Fig. S2**). Additionally, in J4-2, a partial sequence confirms the presence of 1.1 kb ISH11 insertion flanked by a 10 bp direct repeat, while a large 3000+ bp insertion in K3 returned a partial sequence of ISH4. The partial sequence suggests multiple copies of ISH4, or possibly a composite transposon.

**Table 5.**
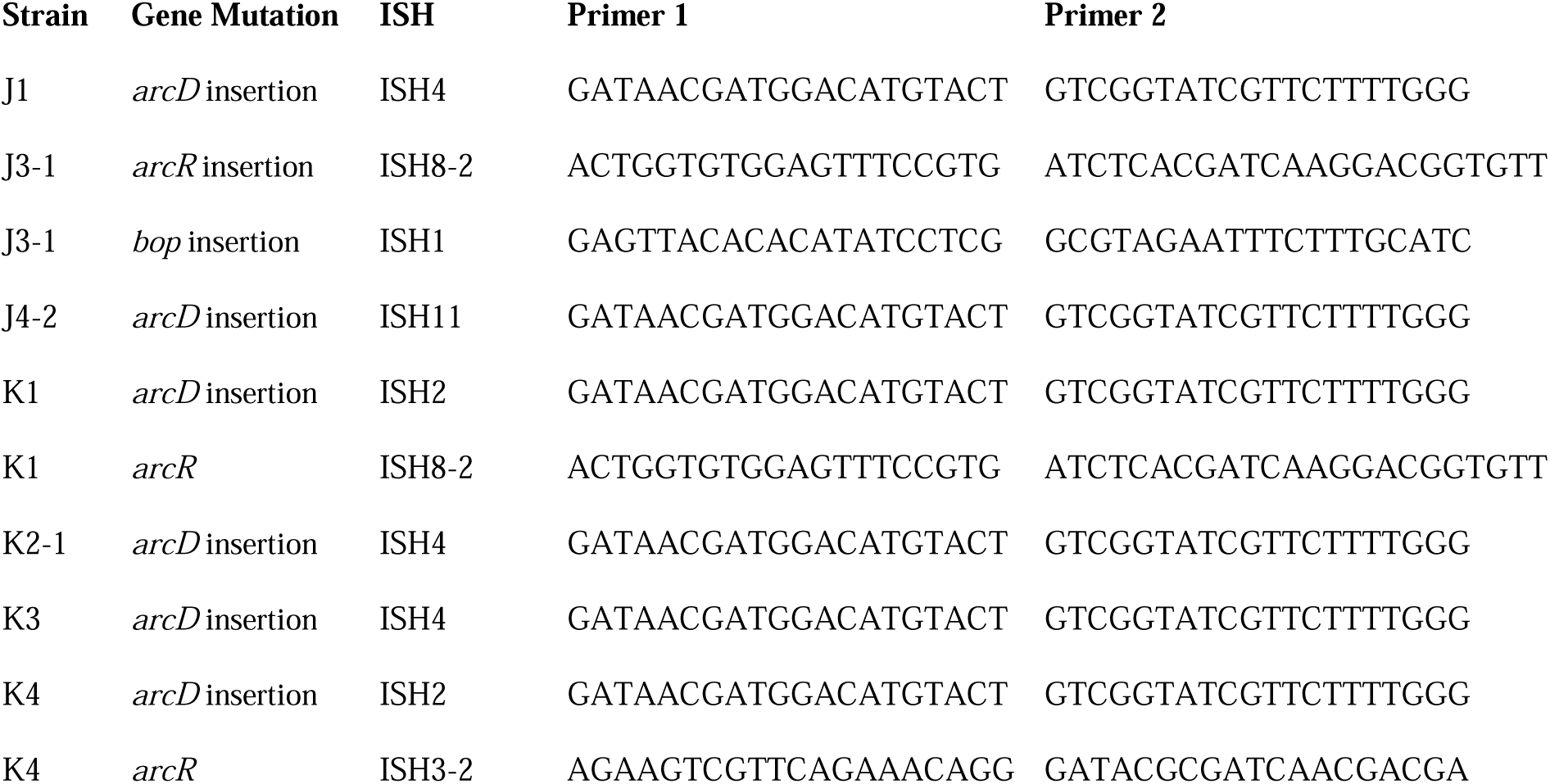
ISH insertions confirmed by PCR in acid-adapted strains.

Four acid-evolved genomes (J-3, K-1, K2-1, K4-1) and one non-acid-evolved clone (M3-1) possess ISH insertions at different sites in *arcR* on pNRC100 (Additional File 1, **Table S3**). ArcR mediates transcriptional regulation of the *arcABDCR* operon for arginine catabolism (40,41). These components include the arginine deiminase (*arcA*), ornithine carbamoyltransferase (*arcB*), carbamate kinase (*arcC*), and the arginine-ornithine antiporter (*arcD*).

For comparison, a remarkably similar system of arginine catabolism reverses acidification for the periodontal bacterium *Streptococcus gordonii* (31,32). Arginine catabolism releases CO_2_ and two molecules of ammonia, which cause net alkalization. The system mediates tooth biofilm formation by *S. gordonii*. For *E. coli*, the arginine decarboxylase Adi reverses acidification at extreme low pH (52). The *adi* system of *E. coli* is induced by acid stress but largely lost by insertion-sequence mutations after long-term evolution in acid (20,22). This suggests a model for acid adaptation in haloarchaea that is remarkably similar to that observed in *E. coli*, in which acid-stress adaptations are knocked down by long-term acid exposure (21).

### Acid-adapted clones shared mutations in bacteriorhodopsin (*bop*)

In NRC-1, our acid-evolved clones J3-1 and K1 each contained an ISH element in the gene *bop* that encodes the light-driven proton pump bacteriorhodopsin (35). The J3-1 allele was confirmed by Sanger sequence as a 1.1 kb insertion of ISH1 with an eight bp target site duplication in *bop* (**Table 5**; Additional File 1, **Fig. S2**). This exact mutation has been previously studied in bacteriorhodopsin mutants, and was in fact the first transposable element identified in haloarchaea (35). This particular target site duplication was shared with acid-evolved clone K1. At a different position, a *bop* ISH insertion was found in one of the M population clones (M3-1) which had not undergone acid selection, consistent with previous spontaneous insertions in this gene.

The *bop* and *arcD* mutations were found together in J3-1, but also in acid-adapted K1, which did not show a significant phenotype under our conditions tested. We inspected strain J3-1 for candidate mutations that might be responsible for this strain’s unique degree of adaptation at low pH. Overall, the J3-1 genome had 16 mutations compared to the NRC-1 ancestor (**Table 6**). Of these, only one mutation affected a gene not affected in any other evolved clone. This is a missense mutation in a ferredoxin gene (*VNG1561*) resulting in a conservative change from lysine to arginine. Mutations were also found affecting several proteins involved in transcriptional regulation, which in combination might contribute to the acid fitness phenotype.

**Table 6.**
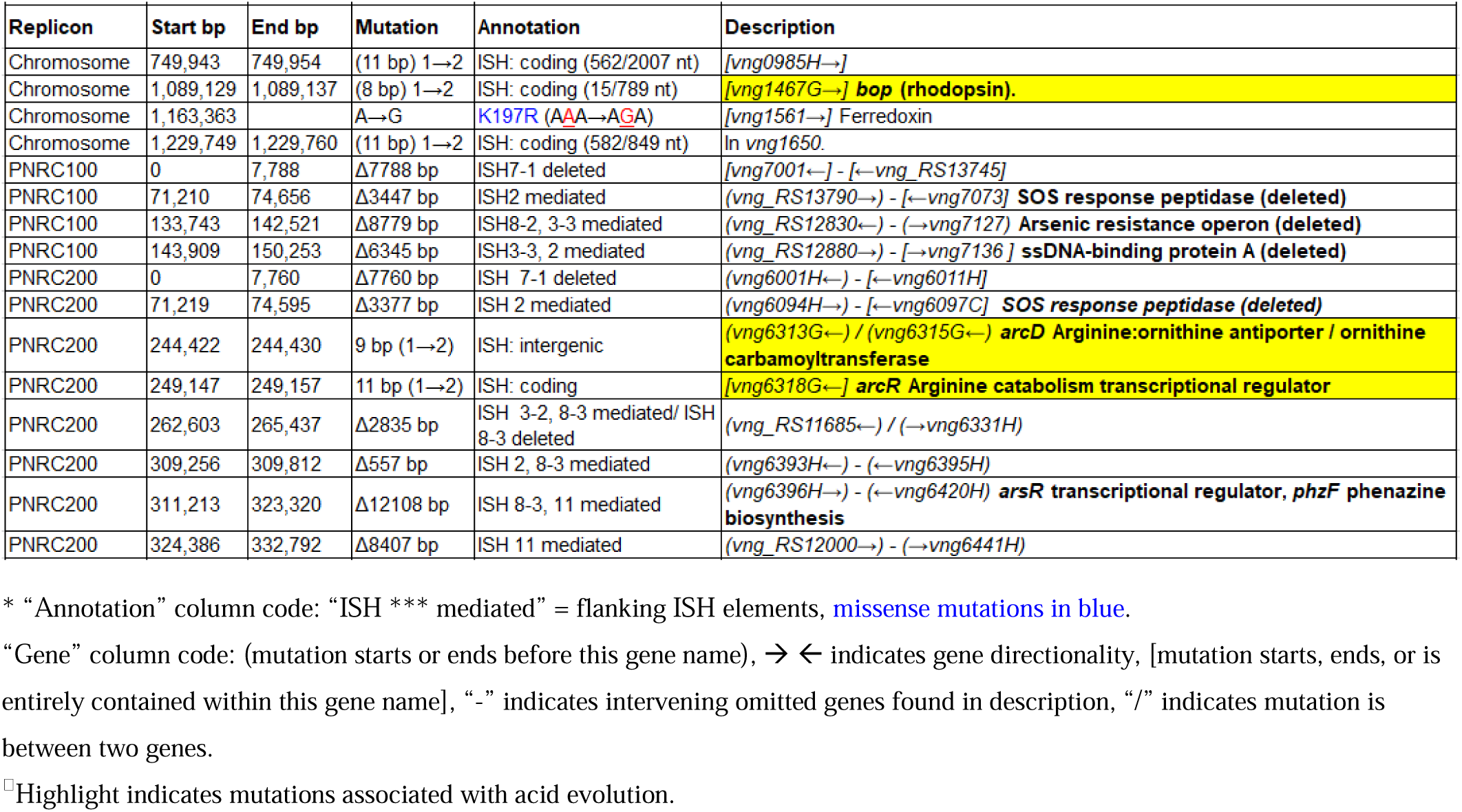
Acid-evolved clone J3-1 mutations.*^□^

| Replicon | Start bp | End bp | Mutation | Annotation | Description |
| --- | --- | --- | --- | --- | --- |
| Chromosome | 749,943 | 749,954 | (11 bp) 1→2 | ISH: coding (562/2007 nt) | [vng0985H→] |
| Chromosome | 1,089,129 | 1,089,137 | (8 bp) 1→2 | ISH: coding (15/789 nt) | [vng1467G→] **bop (rhodopsin).** |
| Chromosome | 1,163,363 |  | A→G | K197R (AAA→AGA) | [vng1561→] Ferredoxin |
| Chromosome | 1,229,749 | 1,229,760 | (11 bp) 1→2 | ISH: coding (582/849 nt) | In vng1650. |
| PNRC100 | 0 | 7,788 | Δ7788 bp | ISH7-1 deleted | [vng7001←] - [←vng_RS13745] |
| PNRC100 | 71,210 | 74,656 | Δ3447 bp | ISH2 mediated | (vng_RS13790→) - [←vng7073] **SOS response peptidase (deleted)** |
| PNRC100 | 133,743 | 142,521 | Δ8779 bp | ISH8-2, 3-3 mediated | (vng_RS12830←) - (→vng7127) **Arsenic resistance operon (deleted)** |
| PNRC100 | 143,909 | 150,253 | Δ6345 bp | ISH3-3, 2 mediated | (vng_RS12880→) - [→vng7136] **ssDNA-binding protein A (deleted)** |
| PNRC200 | 0 | 7,760 | Δ7760 bp | ISH 7-1 deleted | (vng6001H←) - [←vng6011H] |
| PNRC200 | 71,219 | 74,595 | Δ3377 bp | ISH 2 mediated | (vng6094H→) - [←vng6097C] **SOS response peptidase (deleted)** |
| PNRC200 | 244,422 | 244,430 | 9 bp (1→2) | ISH: intergenic | (vng6313G←) / (vng6315G←) **arcD Arginine:ornithine antiporter / ornithine carbamoyltransferase** |
| PNRC200 | 249,147 | 249,157 | 11 bp (1→2) | ISH: coding | [vng6318G←] **arcR Arginine catabolism transcriptional regulator** |
| PNRC200 | 262,603 | 265,437 | Δ2835 bp | ISH 3-2, 8-3 mediated/ ISH 8-3 deleted | (vng_RS11685←) / (→vng6331H) |
| PNRC200 | 309,256 | 309,812 | Δ557 bp | ISH 2, 8-3 mediated | (vng6393H←) - (←vng6395H) |
| PNRC200 | 311,213 | 323,320 | Δ12108 bp | ISH 8-3, 11 mediated | (vng6396H→) - (←vng6420H) **arsR transcriptional regulator, phzF phenazine biosynthesis** |
| PNRC200 | 324,386 | 332,792 | Δ8407 bp | ISH 11 mediated | (vng_RS12000→) - (→vng6441H) |

\* “Annotation” column code: “ISH \*\*\* mediated” = flanking ISH elements, missense mutations in blue.
“Gene” column code: (mutation starts or ends before this gene name), → ← indicates gene directionality, [mutation starts, ends, or is entirely contained within this gene name], “-” indicates intervening omitted genes found in description, “/” indicates mutation is between two genes.
□ Highlight indicates mutations associated with acid evolution.

#### Clones evolved at pH 7.5 show no increase in relative fitness

All evolved clones from generation 500 with Vac^−^ phenotypes were grown over 200 hours in unbuffered CM^+^ medium without acid or iron amendment and compared to the growth phenotype of the NRC-1 Vac^−^ control strain (**Fig. S2A**). Similarly, the growth phenotypes in unstressed medium of Vac^+^ clones from the 500-generation populations that retained them were compared to that of the NRC-1 Vac^+^ ancestor (**Fig. S2B)**. None of the M populations show a significant growth advantage compared to the ancestral strain (Additional File 1, **Fig. S3A and B**).

Growth curves were also conducted for clones from the S populations (evolved with 600 µM FeSO_4_). Media contained CM+ pH 7.5 with 100 mM MOPS and 600 µM FeSO_4_. All evolved clones were persistent Vac^−^ mutants at generation 500 and were therefore compared to an NRC-1 Vac^−^ control (Additional File 1, **Fig. S4**). No significant differences were observed.

#### Multiple clones lost gas vesicles and arsenic resistance

Under laboratory conditions, gas vesicle-producing (Vac^+^) NRC-1 clones have high rates of spontaneous mutation to a vesicle-deficient (Vac^−^) phenotype due to mutations in *gvp* on pNRC100 (38,47). Twelve out of sixteen of our evolved clones, including representatives of each selection type, had lost genes required for gas vesicle nanoparticle production (*gvp*) (53–55). Cultures were oxygenated continually by rotating in a bath, effectively eliminating the competitive advantage of producing gas vesicles in oxygen-limiting environments. Thus, as expected, many insertions and deletions were found that had eliminated gas vesicles (45,47). We characterized gas vesicle phenotypes every 100 generations for the stressed condition populations. These Vac phenotypes (loss of gas vesicle nanoparticles) are presented by population and organized by respective evolution condition in **Table 7**. All evolving populations showed loss of gas vesicle production in some cells. By generation 500, the Vac^-^ phenotype was prevalent in all populations. There was no significant correlation with pH or with iron amendment.

**Table 7:** Change in gas vesicle phenotype during evolution across populations*

| Media condition | Strain | Generation |  |  |  |  |
| --- | --- | --- | --- | --- | --- | --- |
|  |  | 100 | 200 | 300 | 400 | 500 |
| pH 6.3<br>100 mM PIPES | **J1** | + | Vac<sup>+/-</sup> | - | - | - |
|  | **J2** | + | Vac<sup>+/-</sup> | - | - | Vac<sup>+/-</sup> |
|  | **J3** | >1% Vac<sup>-</sup> | Vac<sup>+/-</sup> | - | - | Vac<sup>+/-</sup> |
|  | **J4** | + | + | - | - | Vac<sup>+/-</sup> |
| pH 7.5<br>100 mM MOPS | **M1** | + | + | + | Vac<sup>+/-</sup> | Vac<sup>+/-</sup> |
|  | **M2** | + | + | + | Vac<sup>+/-</sup> | Vac<sup>+/-</sup> |
|  | **M3** | + | + | + | Vac<sup>+/-</sup> | Vac<sup>+/-</sup> |
|  | **M4** | + | >1% Vac<sup>-</sup> | + | + | Vac<sup>+/-</sup> |
| pH 6.3<br>100 mM PIPES<br>600 μM FeSO<sub>4</sub> | **K1** | + | Vac<sup>+/-</sup> | - | Vac<sup>+/-</sup> | Vac<sup>+/-</sup> |
|  | **K2** | Vac<sup>+/-</sup> | Vac<sup>+/-</sup> | - | - | Vac<sup>+/-</sup> |
|  | **K3** | + | Vac<sup>+/-</sup> | - | - | Vac<sup>+/-</sup> |
|  | **K4** | + | Vac<sup>+/-</sup> | - | - | Vac<sup>+/-</sup> |
| pH 7.5<br>100 mM MOPS<br>600 μM FeSO<sub>4</sub> | **S1** | + | + | - | - | - |
|  | **S2** | + | + | - | - | - |
|  | **S3** | + | + | - | - | - |
|  | **S4** | + | + | + | - | - |

\* “+” indicates gas vesicle-forming, “-” indicates non gas vesicle-forming

In addition, 13/16 evolved clones had lost the major arsenic resistance operon (*ars*) encoded on pNRC100 (7). Other mutations affecting transcriptional regulators and initiation factors occurred in parallel across multiple populations. These and other parallel mutations are summarized in **Table 2**. Various hot spots for mutation appear, many of which are caused by ISH insertions or ISH-mediated deletions.

## DISCUSSION

Here we report one of the first evolution experiments to be conducted on a haloarchaeon. A previous evolution experiment involves selection of mutants resistant to ionizing radiation (28).

We compared four environmental conditions: low pH versus optimal pH 7.5, with or without iron supplementation. Overall, in the 500-generation evolved strains, we found a striking pattern of large ISH-mediated deletions, particularly in the two minichromosomes (Additional File 1, **Tables S1-S3**). For comparison, in *E. coli*, experimental evolution for 2,000 generations at low pH yields only occasional large deletions (20,21). However, after just 500 generations of evolution in the haloarchaeon NRC-1, every evolved clone contained several large-scale deletions. ISH insertion mutations greatly outnumbered SNPs. These types of changes reflect frequent DNA rearrangements and genetic variability observed previously in NRC-1 (35,39,49).

The acid-adapted NRC-1 populations showed a striking prevalence of mutations affecting the *arcD* and *arcR* components of arginine transport and catabolism. Arginine catabolism with ammonia release plays a major role in reversing acidity for Gram-positive and Gram-negative bacteria. It is striking to see how the role of acid-dependent arginine catabolism may extend to haloarchaea. The arginase/arginine deiminase family (COG0010) represents a set of orthologs proposed to be among those transferred horizontally to archaea from an ancient bacterial ancestor (56).

The ISH insertions seen in acid-adapted clones would be expected to knock out the arginine system, as seen in *E. coli* experimental evolution with acid (20,41). The reason for the evolutionary loss is proposed to be a readjustment to long-term acid exposure, for which the sustained induction of arginine catabolism becomes counterproductive. It is interesting to find evidence for a similar evolutionary mechanism in a haloarchaeon.

In addition, the acid-evolved strains J3-1 and K1 show an identical insertion mutation affecting the bacteriorhodopsin *bop* gene. The loss of *bop* may be neutral or advantageous under low external pH, where a high proton motive force already exists. The bacteriorhodopsin pump could be a source of proton leakage at high PMF.

The acid-fitness advantage of clone J3-1 could arise from a single mutation unique to J3-1, such as the missense mutation in a ferredoxin that is unique to J3-1. More likely, however, acid fitness arises from a cumulative effect of loss of function mutations in a number of other genes including *arcA, arcR*, and *bop*. It is possible that some unknown factor accounts for the acid-fitness phenotype exhibited by J3-1 under the conditions tested. Nonetheless, it is interesting that the three genes with mutations prevalent in acid-evolved strains all encode products involved in proton consumption or export.

Our findings support previous reports of the importance of ISH elements in haloarchaeal evolution (45), and the observations in *Sulfolobus* that large deletions and loss of function mutations are fitness tradeoffs for surviving in stressful environments (57). Large deletions and IS insertions are also common in experimental evolution of bacteria (20,21,27,58). We also find evidence for accumulation of ploidy changes for the shorter minichromosome, pNRC100 (51). We show that experimental evolution is an effective approach to identify candidate genes for environmental stress response in a haloarchaeon.

## MATERIALS AND METHODS

### *Halobacterium* strains and media

All evolved clones were derived from a stock of *Halobacterium* sp. NRC-1 from the laboratory of Shiladitya DasSarma (3). Liquid cultures were grown in Complex Medium Plus Trace Metals (CM^+^) based on Ref (2), Protocol 25: 250 g/l NaCl, 20 g/l MgSO_4_•7H_2_O, 2 g/l KCl, 3 g/l Na_3_C_6_H_5_O_7_ •2H_2_O, 10 g/l Oxoid Peptone, and 100 μl/l Trace Metals (3.5 g/l FeSO_4_□7H_2_O, 0.88 g/l ZnSO_4_•7H_2_O, 0.66 g/l MnSO_4_•H_2_O, and 0.2 g/l CuSO_4_•5H_2_O dissolved 0.1M HCl) with supplements as needed for the conditions examined (59). CM^+^ solid medium included addition of 20 g/l granulated agar. All cultures were incubated at 42°C with rotation. Cultures on solid media were incubated at 42°C for 7–10 days until colonies reached approximately 1 mm in diameter. A Vac^-^ mutant of our NRC-1 stock culture was obtained by picking a Vac^-^ colony followed by three restreaks on CM^+^ agar.

Liquid CM^+^ media for experimental evolution was made with either 100mM PIPES (pKa=6.8) or 100mM MOPS (pKa=7.2) buffer with pH adjusted using 5 M NaOH or 5 M HCl as needed, followed by filter sterilization. 100 mM FeSO_4_ stock was prepared in deionized water and filter-sterilized before every other dilution during serial batch culture evolution. Sterilized FeSO_4_ stock was added to buffered CM^+^ after filter sterilization. For freezer stocks, live cultures were mixed 1:1 with a 50% glycerol, 50% complex medium basal salts mixture as a cryoprotectant. Complex medium basal salts were 250 g/l NaCl, 20 g/l MgSO_4_•7H_2_O, 2 g/l KCl, 3 g/l Na_3_C_6_H_5_O_7_•2H_2_O. Acidic, control, iron-rich and acidic, and iron-rich media used in the evolution consisted of: CM^+^ pH 6.5 with 100 mM PIPES (populations J1-J4), CM^+^ pH 7.5 with 100 mM MOPS (populations M1-M4), CM^+^ pH 6.5 (or pH 6.3) with 100 mM PIPES 600 µM FeSO_4_ (populations K1-K4), and CM^+^ pH 7.5 with 100 mM MOPS 600 µM FeSO_4_ (populations S1-S4).

### Experimental evolution

A total of 16 populations (four per evolution condition) were founded from a 5 ml CM^+^ tube culture (7-10 days incubation) of *Halobacterium* sp. NRC-1 that was diluted 500-fold and incubated 4 days in a 42°C shaker bath at 200 rpm. At the end of the fourth day, 10 µl of the previous culture was diluted into 5 ml of fresh CM^+^ media amended as necessary for the respective stress condition. The resulting 1:500 dilutions yield approximately nine generations per dilution cycle. If cultures did not reach a healthy cell density as qualitatively evaluated for each dilution, 1:100 or 1:250 dilutions were performed to prevent loss of evolving populations. Alternative dilution concentrations were factored into total generation counts at the end of experimental evolution. When evolution was interrupted, the populations were revived by 1:250 dilutions from freezer stocks of the previous dilution. Freezer stocks comprised 1 ml liquid, mature haloarchaea culture for each evolving population and 0.5 ml glycerol/basal salts mixture, stored in 2 ml Wheaton brand vials and frozen at −80°C for each dilution, totaling 16 freezer stocks every four days. A summary of the evolution procedure is presented in **Figure S1**.

### Clone selection

Clones were isolated by plating 10 µl of culture from generation 100, 200, 300, 400, and 500 from freezer stocks for all 16 evolving populations on CM^+^ agar plates, followed by incubation in a sealed container at 42°C for 7–10 days. Isolated colonies were then selected for diverse Vac phenotypes, streaked on fresh CM^+^ agar plates, and incubated a second time. The process was repeated a third time to ensure isolation of select genetically pure clones. Colonies from the third streak were grown in unbuffered CM^+^ pH 7.2, and stocks were frozen for later phenotype and genotype characterization. One clone was isolated from each population every 100 generations. For populations that presented mixed gas vesicle production phenotypes, we isolated both a Vac^+^ clone and a Vac^-^ clone. In total, 75 clones were isolated from generation 100, 200, 300, and 400 of the evolution. Clones were similarly isolated from generation 500; however, the first streak was taken directly from evolving populations, rather than from frozen stock in Wheaton vials. Two clones were isolated from each population at 500 generations, for a total of 32 clones.

### Gas vesicle formation phenotype analysis

Vesicle formation phenotype was assessed qualitatively based on the relative translucence of plated colonies and denoted as Vac^+^ or Vac^−^ as appropriate (2,47). If more than one Vac phenotype was observed in a streak during strain isolation, the phenotypic variant colonies were re-streaked and treated as separate clonal isolates. Vac phenotypes were evaluated for persistence with each streak based on whether or not Vac^+^ colonies yielded >1% Vac^−^ progeny or vice versa.

### Growth assays

The generation 500 clones used in these assays are summarized in **Table 1**. Clones were cultured in unbuffered CM+ at pH 7.2, and incubated for four days in a 42°C shaker bath with 200 rpm orbital aeration. Over-week starter cultures were diluted 1:1000 into new test tubes with 5 ml of the appropriate test condition media. A media blank was included for each media condition, and each clone was tested with four to eight biological replicates, depending on the assay. Immediately after inoculation, OD_600_ values were recorded by a Spectramax 384+ spectrophotometer at 600 nm using Softmax Pro version 6.4.2. Daily readings were taken for nine days. Media for these tests included CM+ pH 6.3 100 mM PIPES and CM+ pH 6.1 100 mM PIPES for J clones. M clones were tested in CM+ pH 7.5 100 mM MOPS. K clones were tested in CM+ pH 6.3 100 mM PIPES 600 µM FeSO_4_ and CM+ pH 6.1 100 mM PIPES 600 µM FeSO_4_. S clones were tested in CM+ pH 7.5 100 mM MOPS 600 µM FeSO_4_.

To test for pH-dependent growth advantages, evolved clones that showed growth advantages over ancestor in their respective evolution stress conditions under which they were evolved were also tested for growth advantages in pH conditions other than those in which they evolved. For these experiments, J3-1 was cultured in CM+ pH 7.5 100 mM MOPS and compared using a Vac^+^ NRC-1 control, M3-1 was cultured in CM+ pH 6.1 100 mM PIPES and compared using a Vac^+^ NRC-1 control, and K2-1 was cultured in CM+ pH 7.5 100 mM MOPS 600 µM FeSO_4_ and compared to both Vac^+^ and Vac^−^ NRC-1 controls due to gas vesicle phenotype ambiguity. Analysis was carried out with comparisons to an ancestral control expressing the same Vac phenotype as the evolved clone.

All growth assays were evaluated for statistical significance using ANOVA test with Tukey post-hoc or paired T-test using base R and agricolae package. Comparisons between clones were made using post log-phase endpoint “E” values for optical density at six days post inoculation.

### DNA extraction and genome sequencing

Genomic DNA was isolated from the 16 evolved clones and the ancestor NRC-1 using an Epicentre MasterPure Gram Positive DNA Extraction Kit and a modified procedure. Lysozyme was omitted, and DNA purity and concentration was determined using a Thermo Scientific NanoDrop 2000. Genomic DNA was sequenced at the Michigan State University Research Technology Support Facility (RTSF) Genomics Core. Libraries were prepared using the Illumina TruSeq Nano DNA library preparation kit for Illumina MiSeq sequencing and loaded on a MiSeq flow cell after library validation and quantitation. Sequencing was completed using a 2-by 250-bp paired-end format using Illumina 500 cycle V2 reagent cartridge. Illumina Real Time Analysis (RTA) v1.18.54 performed base calling, and the output of the RTA was demultiplexed and converted to FastQ format with Illumina Bcl2fastq v1.8.4.

### Sequence assembly and analysis using the *breseq* computational pipeline

The computational pipeline *breseq* version 0.27.1 was used to assemble and annotate the resulting Illumina reads of the evolved clones (42–44). The current *breseq* version is optimized to detect IS element insertions and IS-mediated deletions, as well as SNPs and other mutations in *E. coli* (19). Illumina reads were mapped to the *Halobacterium* sp. NRC-1 reference genome (NCBI GenBank assembly accession GCA_000006805.1). Mutations were predicted by *breseq* through sequence comparisons between the evolved and ancestral clones.

The Integrative Genomics Viewer (IGV) from the Broad Institute at Massachusetts Institute of Technology was used to visualize the assembly and mutations in the evolved clonal sequences mapped to the reference NRC-1 genome (60). Each replicon was mapped separately using the following RefSeq IDs: NC_002607.1 (main chromosome), NC_001869.1 (pNRC100), and NC_002608.1 (pNRC200). Sequence mean coverage in each evolved clone was estimated using the *breseq* fit dispersion function.

### PCR confirmation of ISH insertions

PCR primers (Table 5) were designed to confirm the presence of insertion sequences at hypothetical target site duplications. Primers adhered to the following specifications using Sigma Aldrich Oligo Evaluator: 19-22 bp in length, GC content between 40-60%, no single bp runs >3, weak to no secondary structure, and no primer dimer. Oligos were checked for sequence identity of ≤13 bp to any part of the NRC-1 genome other than the target site using NCBI BLAST. We ran 50-µl PCR using Applied Biosystems Amplitaq Gold 360 Master Mix according to the package insert with 50 µl reaction containing GC enhancer. To assess insert length, 10 µl of PCR product was electrophoresed in a 1% agarose gel. PCR products were then purified either by Qiagen QIAquick PCR Purification Kit or QIAquick Gel Extraction Kit.

### Accession number for sequenced genomes

Sequenced genomes are deposited under SRA accession number SRP195828.

## ACKNOWLEDGEMENTS

This project was supported by the National Science Foundation award MCB-1613278 to Joan Slonczewski; by the 2016 NASA Astrobiology Program Early Career Collaboration Award to Karina Kunka and Jessie Griffith; and by NASA Exobiology grants NNX15AM07G and NNH18ZDA001N to Shiladitya DasSarma. We thank Friedhelm Pfeiffer for pointing out the *arcD* annotation. We thank Landon Porter for expert technical assistance.

